# Structural Mechanism of Specific Nucleobase Recognition by a Monoclonal Antibody in CoolMPS™ Sequencing

**DOI:** 10.64898/2025.12.22.696117

**Authors:** Lixiang Yang, Qiyue Wang, Hanyi Liao, Weiwei Li, Chang Xu, Jianxun Qi, Huijun Zhang, Lifeng Fu, Meng Yang

**Affiliations:** MGI Tech, Shenzhen 518083, China; Key Laboratory of Pathogen Microbiology and Immunology, Institute of Microbiology, Chinese Academy of Sciences, Beijing 100101, China; Medical School, University of Chinese Academy of Sciences, Beijing 101408, China; Faculty of Health Sciences, University of Macau, Macau, SAR, 999078, China

## Abstract

Massively parallel sequencing (MPS) has revolutionized genomics, yet traditional methods utilizing fluorescently labeled nucleotides suffer from “scarring” effects that limit read accuracy. The CoolMPS™ technology overcomes this by using unlabeled reversible terminators (RTs) with a 3’-O-azidomethyl blocking group, detected by highly specific fluorescent antibodies. However, the atomic-level mechanism by which these antibodies discriminate between modified and natural nucleotides within a DNA strand remains unclear. Here, we report the crystal structures of the “A-fab” antibody fragment in complex with the monomeric antigen 3’-O-azidomethyl-dATP (dATP-N_3_) at 2.50 Å resolution, and with an one strand of the duplex carries an azide-modified deoxyadenosine nucleotide at its 3′ terminus (dsDNA-dATP-N_3_) at 1.99 Å resolution. Isothermal titration calorimetry (ITC) revealed that A-fab binds dATP-N3 with nanomolar affinity (*K*_*D*_≈17.14 nM), while binding to natural dATP is negligible (*K*_*D*_≈7.51µM), representing a ∼440-fold specificity. Structural analysis reveals a conserved hydrophobic pocket formed by CDR residues (e.g., Trp79, Tyr53, Leu67) that specifically accommodates the 3’-azidomethyl group. This interaction is critical for affinity and is maintained in the dsDNA-ATP-N3 complex, validating the antibody’s function in a sequencing context. These findings provide the structural rationale for the high fidelity of the CoolMPS™ platform.

## 1. Introduction

The rapid evolution of Next-Generation Sequencing (NGS) has fundamentally transformed biological research and clinical diagnostics [1, 2]. The dominant Sequencing-by-Synthesis (SBS) chemistry typically relies on dye-labeled reversible terminators (RTs), where a fluorophore is attached to the nucleobase via a cleavable linker [3, 4]. While successful, this approach faces inherent limitations: incomplete cleavage of the linker leaves a chemical “scar” on the nucleobase. Accumulation of these scars over sequencing cycles can distort the DNA double helix, inhibit polymerase activity, and lead to phasing errors, thereby limiting read lengths and quality [5, 6].

To address these challenges, “Cold” MPS (CoolMPS™) technology was developed, utilizing unlabeled RTs with a 3’-O-azidomethyl blocking group [7]. Instead of direct laser excitation of the nucleotide, base detection is achieved via fluorophore-labeled antibodies that specifically bind to the incorporated RTs [7]. This method utilizes natural nucleobases without permanent modifications, potentially enabling longer and more accurate reads. The viability of this platform depends entirely on the availability of antibodies that can distinguish the 3’-modified nucleotide from natural DNA with exceptional specificity [8].

Antibodies binding to nucleic acids (anti-DNA antibodies) have been studied in the context of autoimmune diseases and DNA damage recognition [9, 10]. For instance, antibodies against UV-induced DNA photoproducts often employ an “induced-fit” mechanism, where complementarity-determining regions (CDRs) undergo conformational changes to sequester the damaged base [11-13]. However, recognizing a single chemical modification (the 3’-azidomethyl group) on a specific nucleobase (Adenine) presents a unique challenge in molecular recognition. The antibody must possess a “reading head” capable of sensing the modification while excluding natural nucleotides to prevent background noise [14].

In this study, we elucidate the recognition mechanism of “A-fab,” a key component of the CoolMPS™ chemistry. We determined the crystal structures of A-fab in complex with both the free nucleotide (dATP-N_3_) and an dsDNA-dATP-N_3_ containing the modification. Combined with thermodynamic analysis, our results demonstrate how A-fab achieves exquisite specificity through a distinct hydrophobic sub-pocket, providing a structural blueprint for high-accuracy sequencing reagents.

## 2. Materials and Methods

### 2.1 Preparation of modified DNA antigens

The 3’-O-azidomethyl-dATP (dATP-N_3_) used in crystallization experiments was obtained from BGI Research (Wuhan Huada Zhizao Biological Engineering Co., Ltd., Wuhan, China).

High-purity, base-specifically modified double-stranded DNA antigens were prepared using a self-developed enzymatic synthesis protocol under mild reaction conditions. In these antigens, one strand of the duplex carries an azide-modified deoxyadenosine nucleotide at its 3′ terminus (dsDNA-dATP-N_3_). Each dsDNA antigen consisted of a biotinylated duplex in which the 5′ end of one strand was labeled with biotin and the 3′ terminus of the complementary strand carried an azide-blocked deoxynucleotide corresponding to A, T, C, or G. DNA synthesis was catalyzed by a proprietary polymerase system based on the MGISEQ-2000RS FCL PE150 platform (Cat. No. 940-000032-00). Two complementary dsDNA-ATP-N3 were designed for each antigen (Table 2): a full-length template strand (Strand 1, length n) and a shorter complementary primer strand (Strand 2, length n – x, where x is an integer between 1 and n – 5). Strand 2 was synthesized with a 5′ biotin modification to enable subsequent immobilization. For duplex formation, Strand 1 and Strand 2 were mixed at equimolar concentrations (typically 10 μL of 1 μM Strand 1 with 10 μL of 1 μM Strand 2) in annealing buffer. The mixture was heated to 80–95 °C for 5–15 min to denature secondary structures and then gradually cooled to room temperature to allow annealing and formation of the primer–template complex. The annealed dsDNA served as the substrate for the enzymatic extension reaction. Base-specifically modified antigens were generated by extending Strand 2 in the presence of azide-capped deoxynucleotides. The annealed primer–template complexes were incubated with the MGISEQ-2000RS FCL PE150 DNA polymerase system and a single type of modified deoxynucleotide triphosphate (dNTP-N_3_; A, T, C, or G). Briefly, reactions were carried out at 40–50 °C for 30 min to allow incorporation of the azide-capped nucleotide at the 3′ end of Strand 2. By selectively supplying only one type of azide-modified nucleotide in each reaction, four distinct dsDNA antigens—A-, T-, C-, or G-terminated—were directionally synthesized.

### 2.2 Protein Expression and Purification

The coding sequence of A-fab, whose amino-acid sequence was derived from the antibody disclosed in the Chinese patent application CN 113272448 A (MGI)[15], was cloned into a pCAGGS vector containing a His_6_ tag at the C-terminus and transiently transfected into Expi293F™ cells. After 5-day culturing, the supernatant was collected by centrifugation (8000 rpm, 90 min, 4°C), and the soluble protein was purified using Ni affinity chromatography with a HisTrap™ HP 5 mL column (GE Healthcare). Further purification was carried out by size-exclusion chromatography using a Superdex™ 200 Increase 10/300 GL column (GE Healthcare). The proteins were in PBS buffer during purification. Finally, SDS-PAGE was performed to analyze the size and purity of the eluted peak proteins. The purified proteins were concentrated to 10 mg/mL and stored at −80°C.

### 2.3 Crystallization and Data Collection

After purification, the purified A fab protein was concentrated to 5 mg/mL and 10 mg/mL. Crystallization was performed by the sitting-drop vapor diffusion method at 18°C. Purified A fab crystals grew in drops containing 0.2 M Ammonium chloride and 20% w/v PEG 3350 after two weeks. For ligand soaking, crystals were transferred into the reservoir solution supplemented with 30 μM dATP-N3 or 100 μM dsDNA-ATP-N3 and incubated for 4 hours at room temperature. Complex crystals were flash-frozen in liquid nitrogen and sent to Shanghai Synchrotron Radiation Facility (SSRF) for data collection.

### 2.4 Structure Determination and Refinement

Diffraction data were processed using HKL2000. The structures were solved by molecular replacement using Phaser. Refinement was carried out using Phenix and Refmac5, and model building was performed in WinCoot. Data collection and refinement statistics are summarized in Table 1.

**Table 1.**
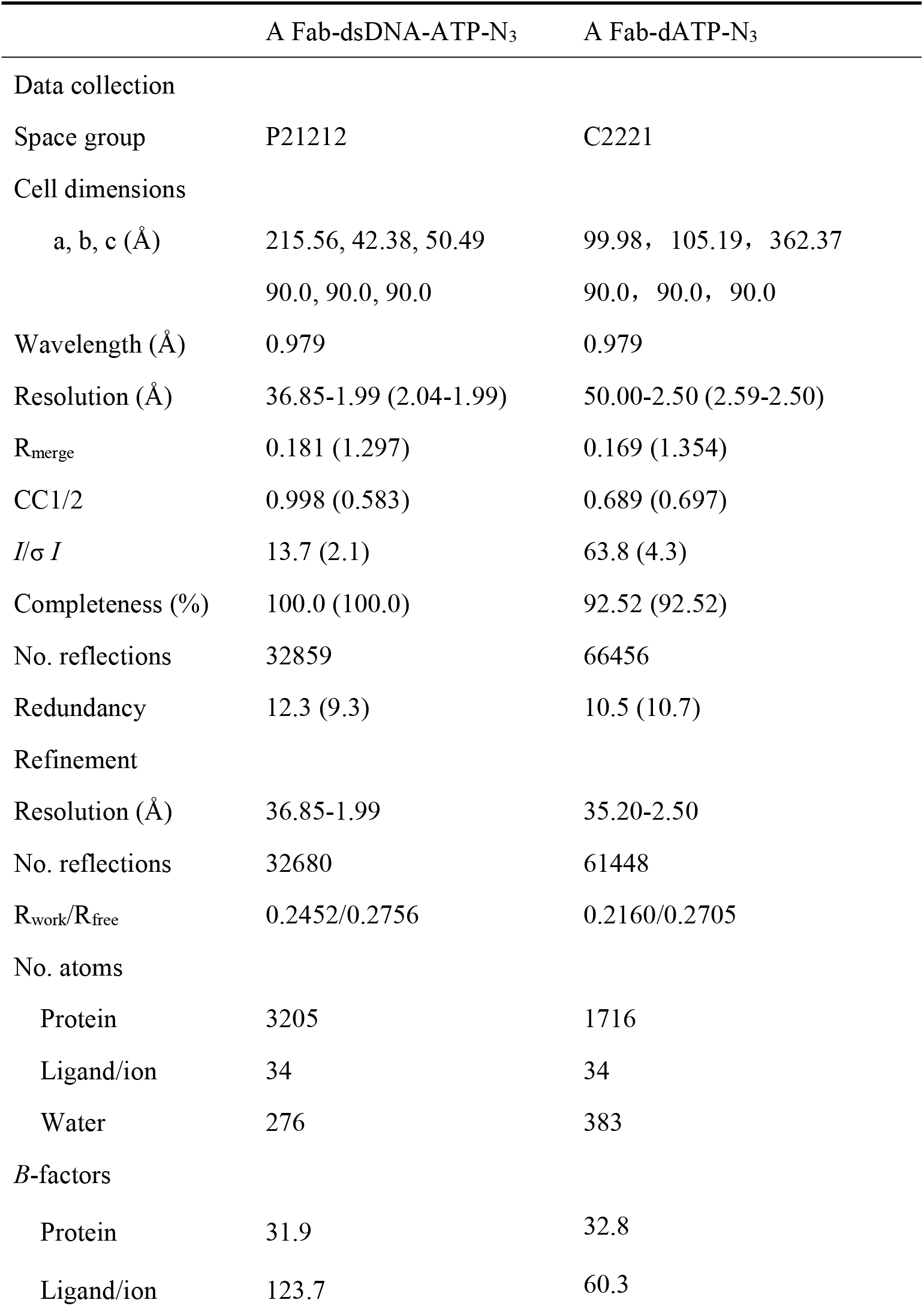
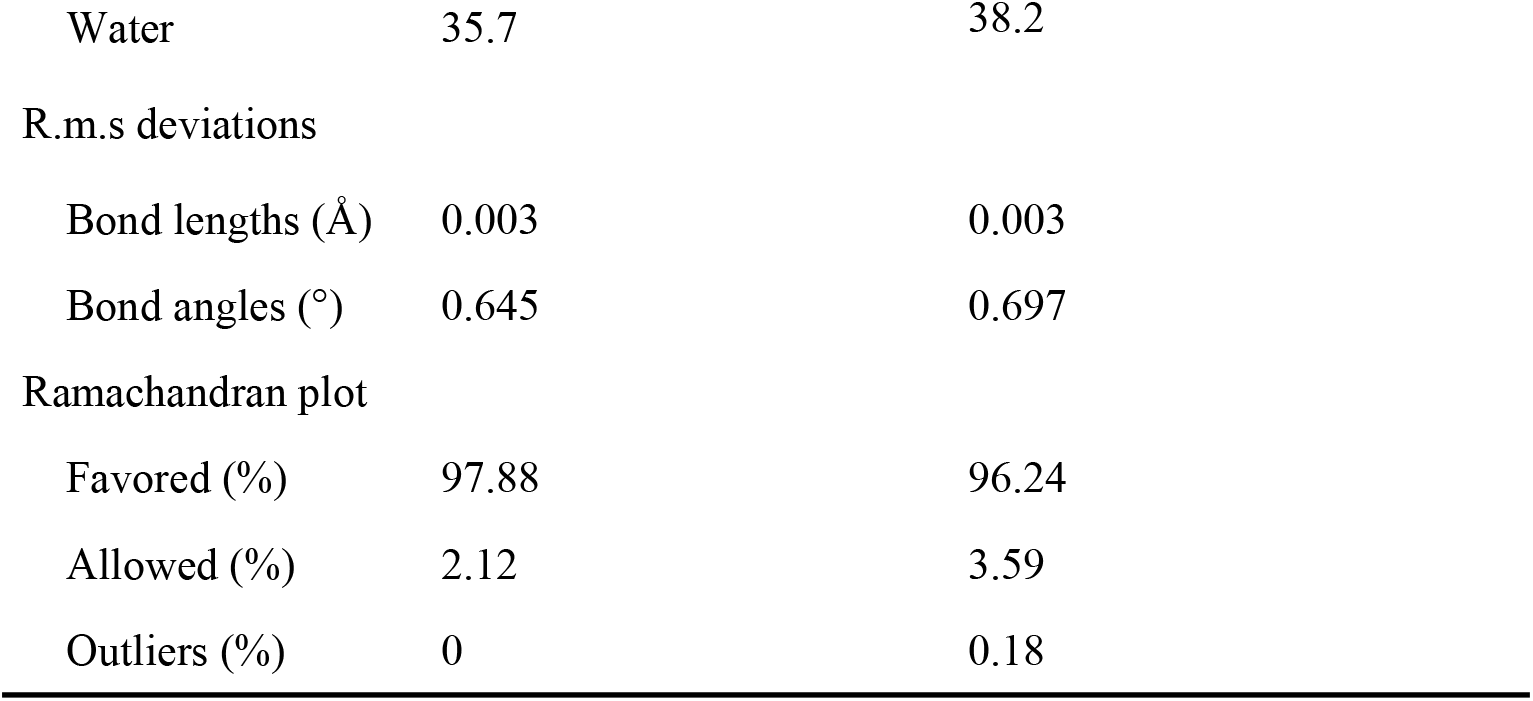
Crystallographic data collection and refinement statistics.

### 2.5 Isothermal Titration Calorimetry (ITC)

Affinity measurements were performed using an Affinity ITC system. The cell contained A-fab (20 µM) in PBS, and the syringe contained either dATP-N_3_ (100 µM) or dATP (200 µM). Titrations were performed at 25°C. Data were fitted to a one-site binding model to determine the dissociation constant (*K*_*D*_).

## 3. Results

### 3.1 Crystallographic Analysis of A-fab Complexes

We successfully determined the crystal structures of A-fab in two states: complexed with the monomeric ligand dATP-N_3_ and complexed with an oligonucleotide (dsDNA-ATP-N_3_) (Table 1). X-ray diffraction data for both complexes were collected at the SSRF at a wavelength of 0.979 Å. The crystals of the Afab–oligonucleotide complex belonged to space group P2_1_2_1_2 with unit cell parameters a = 215.56 Å, b = 42.38 Å, and c = 50.49 Å, diffracting to a maximum resolution of 1.99 Å. This dataset exhibited 100.0% overall completeness (100.0% in the highest shell), with an R_merge of 0.181 (1.297 in the highest shell), CC1/2 of 0.998 (0.583), mean I/σ(I) of 13.7 (2.1), and redundancy of 12.3 (9.3), indicating high data redundancy and a favorable signal-to-noise ratio. The Afab–dATP-N_3_ complex crystallized in space group C222_1_ with unit cell parameters a = 99.98 Å, b = 105.19 Å, and c = 362.37 Å, at a resolution of 2.50 Å. For this dataset, the overall R_merge was 0.169 (1.354), CC1/2 was 0.689 (0.697), mean I/σ(I) was 63.8 (4.3), redundancy was 10.5 (10.7), and completeness in the highest shell was 92.52%, also confirming reliable data quality. Following structural refinement, the Rwork/Rfrevalues were 0.2452/0.2756 for the Afab–oligonucleotide complex (1.99 Å) and 0.2160/0.2705 for the Afab–dATP-N_3_ complex (2.50 Å) (Table 1). The final model of the former comprises 3,205 protein atoms, 34 ligand/ion atoms, and 276 water molecules, with mean B-factors of 31.9 Å^2^ (protein), 123.7 Å^2^ (ligand/ions), and 35.7 Å^2^ (water). The latter model consists of 1,716 protein atoms, 34 ligand/ion atoms, and 383 water molecules, with mean B-factors of 32.8 Å^2^, 60.3 Å^2^, and 38.2 Å^2^, respectively. Ramachandran plot analysis indicated that for the Afab–oligonucleotide structure, 97.88% of residues are in favored regions and 2.12% in allowed regions, with no outliers. For the Afab–dATP-N_3_ structure, 96.24% of residues are in favored regions, 3.59% in allowed regions, and only 0.18% are outliers. Both structures exhibit root-mean-square deviations (RMSD) from ideal geometry of 0.003 Å for bond lengths and approximately 0.65–0.70° for bond angles. Collectively, these statistics demonstrate the high quality of both Afab complex structures, allowing for a detailed elucidation of their ligand-binding modes.

### 3.2 Structural Basis of Specificity for the 3’-O-azidomethyl Group

The high-resolution crystal structures (2.50 Å and 1.99 Å) allow for a precise mapping of the interactions that confer A-fab its exquisite specificity. The crystal structures reveal that the exquisite specificity of A-fab arises from a highly specialized antigen-binding cleft located at the interface of the variable heavy (VH) and variable light (VL) domains. As visualized in the electrostatic surface potential maps (Fig. 1b and 1e), the binding site comprises a positively charged rim that anchors the triphosphate tail, leading into a deep, hydrophobic cavity that accommodates the nucleobase and ribose moieties.

**Figure 1.**
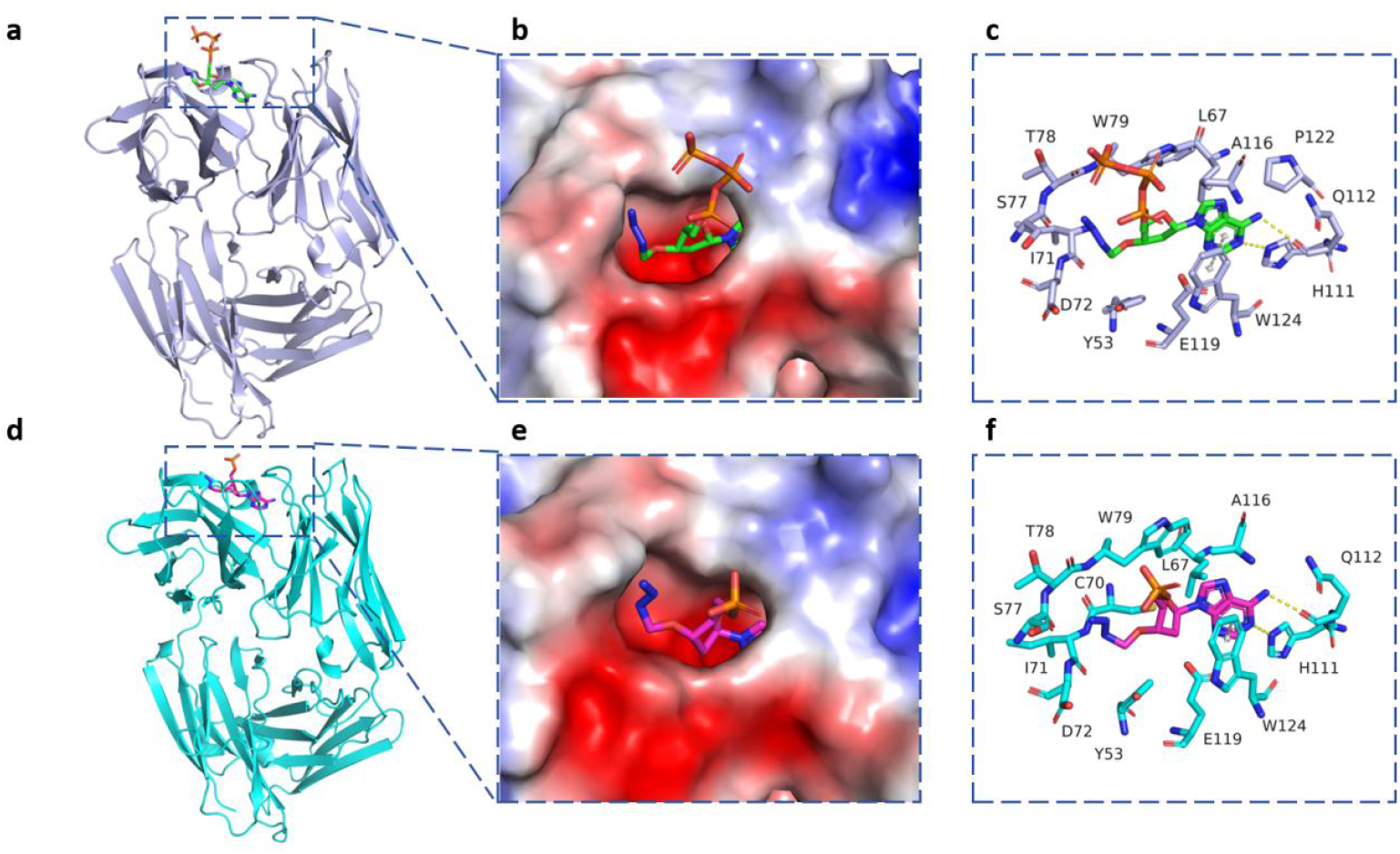
Atomic details of specific recognition by A-fab. (a, d) Overall structures of A-fab in complex with dATP-N_3_ (blue) and dsDNA-dATP-N_3_ (cyan). (b, e) Electrostatic surface potential views of the binding pocket. The surface is colored by potential (blue: positive; red: negative). Note the deep, shape-complementary cavity that engulfs the 3’-azidomethyl group (center), while the phosphate groups interact with the electropositive rim (blue region). (c, f) Detailed view of the antigen-binding site. The 3’-O-azidomethyl group is sequestered in a hydrophobic cage formed by the “lid” residue Tyr53, Trp79, flanked by Leu67 and Ala116. The adenine base is stabilized by π-π stacking with Trp124 and hydrogen bonds (yellow dashes) with His111/Gln112. The structural arrangement is conserved between the monomeric (c) and dsDNA-dATP-N_3_ (f) complexes.

The atomic details of this recognition are elucidated in Fig. 1c. The adenine base is stabilized by a combination of π-π stacking interactions with Trp124 and specific hydrogen bond networks involving His111 and Gln112, which read the Watson-Crick edge of the base, ensuring base-specific selection.

Most critically, the structure unveils the mechanism for recognizing the 3’-O-azidomethyl modification. The modified 3’-group is not exposed to the solvent but is buried within a dedicated hydrophobic sub-pocket defined by residues Tyr53, Cys70, Trp79, Leu67, Ala116, and Ile71 (Fig. 1c). This pocket forms a “hydrophobic cage” that exhibits perfect shape complementarity to the azidomethyl moiety. The close van der Waals contacts provided by these residues (particularly the indole ring of Trp79, aromatic ring of Tyr53 and the side chain of Leu67) act as a structural “lock.” In the absence of the azidomethyl group (as in natural dATP), this sub-pocket would remain void, leading to a significant loss of packing energy and stabilizing interactions.

To validate whether the antibody maintains this recognition mode during active sequencing, we solved the structure of A-fab in complex with an dsDNA-ATP-N3 containing the incorporated dATP-N3 (Fig. 1d). Comparison of the monomeric complex (Fig. 1a-c) with the dsDNA-ATP-N3 complex (Fig. 1d-f) reveals a remarkable structural convergence.

The overall root-mean-square deviation (RMSD) between the two antibody structures is negligible, indicating a rigid-body binding mode. In the dsDNA-ATP-N3 complex, the phosphate backbone is stabilized by Ser77, Thr78, and Tyr53 (Fig. 1f), residues that are positioned identically to those in the monomeric structure. Crucially, the configuration of the hydrophobic sub-pocket (Trp79, Leu67, Ala116) remains invariant. This confirms that A-fab recognizes the modified nucleotide in a conformation that is accessible within the DNA double helix, without requiring significant distortion of the DNA template or induced-fit rearrangement of the antibody CDRs.

### 3.3 Thermodynamic Validation of Specificity

ITC experiments provided quantitative validation of the structural observations (Figure 2). A-fab bound to dATP-N_3_ with high affinity (*K*_*D*_=17.14±5.64 nM) and a favorable enthalpy change (Fig. 2, left), indicating tight, specific binding. In contrast, binding to natural dATP was drastically weaker (*K*_*D*_=7.51±3.97µM) (Fig. 2, right). The ∼440-fold difference in affinity demonstrates that the antibody relies heavily on the 3’-azidomethyl group for stable complex formation, ensuring low error rates during the CoolMPS sequencing process.

**Figure 2.**
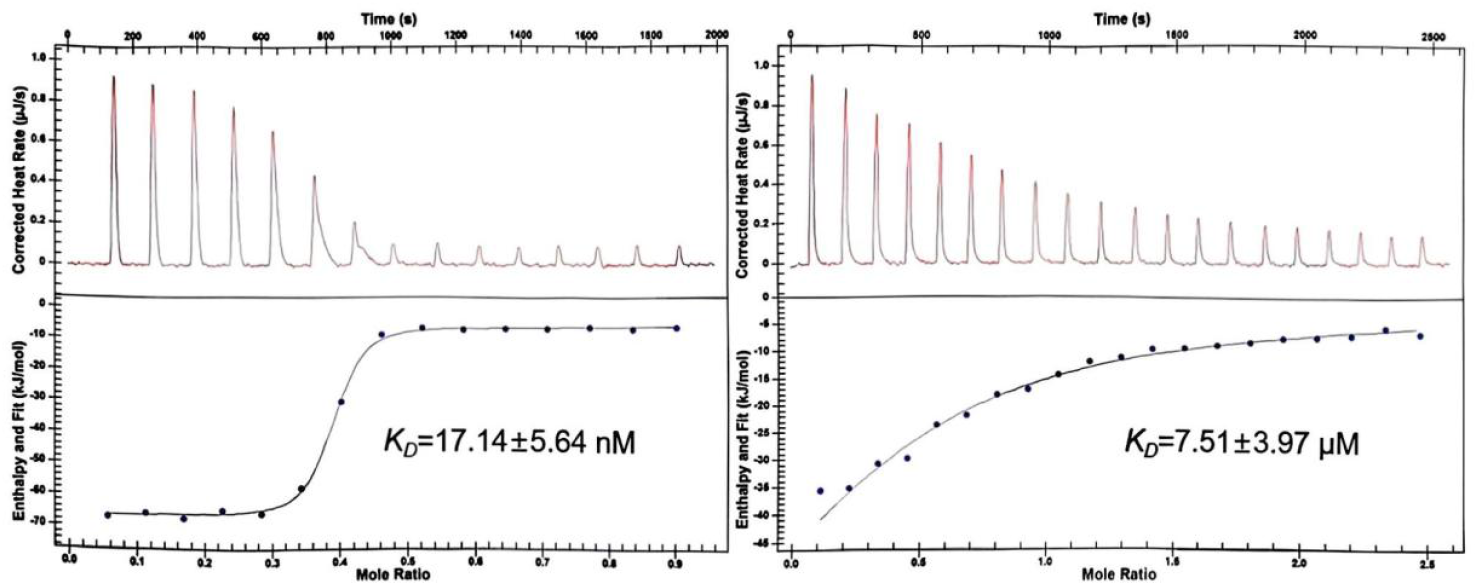
ITC analysis of A-fab binding to dATP-N_3_ and unmodified dATP. Isothermal titration calorimetry (ITC) experiments were performed by titrating dATP-N_3_ (left) or dATP (right) into A-fab at 25 °C. Upper panels show the raw heats of injection (red, integrated peaks; black, fitted baseline) plotted as corrected heat rate versus time. Lower panels show the integrated heats per injection plotted against the molar ratio of ligand to protein, together with the best-fit binding isotherms (solid lines).

## 4. Discussion

Our structural and thermodynamic data validate the CoolMPS™ platform by revealing how A-fab acts as a precise molecular sensor. Specificity is achieved via a “lock-and-key” mechanism where a pre-formed hydrophobic pocket accommodates the 3’-O-azidomethyl group. This interaction drives high-affinity binding to dATP-N_3_ while imposing a severe desolvation penalty on natural dATP, ensuring robust discrimination. The antibody employs a rigid-body binding mode that facilitates rapid kinetics without distorting the DNA backbone, preserving template integrity for subsequent sequencing cycles. Ultimately, A-fab functions as a “molecular caliper” that separates base recognition (via hydrogen bonds and stacking) from modification sensing (via a specialized hydrophobic pocket), minimizing steric mismatches and ensuring high fidelity. These structural insights not only validate the CoolMPS™ chemistry but also offer a rational blueprint for the development of future antibodies targeting novel nucleic acid modifications.

## Data availability

The atomic coordinates and structure factors for the two A Fab–ligand complexes have been deposited with the Worldwide Protein Data Bank (wwPDB). Accession codes will be provided upon release.

## Acknowledgements

This research is supported by Ministry of Science and Technology of the People’s Republic of China’s program titled ‘National Key Research and Development Program of China’ (No. 2022YFF1202200, 2022YFF1202203) and Science, Technology, Innovation Commission of Shenzhen Municipality under grant No. JSGGZD20220822095802006, the Youth Innovation Promotion Association CAS (2023096).

## Author contribution

M.Y. and L.F. conceived and designed the study. L.X.Y. and L.F. performed analysis and wrote the manuscript. L.F. contributed to the design and guidance of the experimental section. Q.W. and H.L. performed modified DNA antigens preparation, protein expression and purification and crystallization. J.Q. collected the structural data and solved the structures. W.L. conducted ITC assay. M.Y. provided suggestions for the writing and revision of the manuscript. All authors read and approved the final version of the manuscript.

## Notes

### Competing Interest Statement

The authors have declared no competing interest.

